# Phenology of *Hyalomma marginatum*: a longitudinal study in a site of southern France

**DOI:** 10.1101/2025.06.30.662279

**Authors:** Frédéric Stachurski, Maxime Duhayon, Anaïs Appelgren, Gilles Balança, Carla Giupponi, Thomas Pollet, Laurence Vial

## Abstract

*Hyalomma marginatum* tick establishment has been recorded in southern mainland France for around ten years. This two-host species is one of the main vectors of the Crimean-Congo haemorrhagic fever (CCHF) virus. Knowing the phenology of the tick is a prerequisite for a better understanding of its spread and of CCHF virus dispersal. A longitudinal study was carried out between 2016 and 2022 in a horse herd to determine the seasonal variations of *H. marginatum* infestation. In the study site, adult ticks started infesting their hosts from the beginning of March. Infestation peaked during the first weeks of May. At that period, on average, up to 4 or 5 ticks attached daily to each horse. Then, the infestation steadily decreased until the end of August. In this site, birds are parasitised by *H. marginatum* immatures (larvae and nymphs) in summer, from mid/late June to the end of September. The larvae are therefore not yet active when the migratory birds pass through in spring. Blackbird, *Turdus merula*, was the most heavily infested bird species: nearly two-thirds of the blackbirds examined in summer were parasitised by *H. marginatum*. It is suggested that this bird species could be a good sentinel host for monitoring the establishment of the tick in currently *H. marginatum*-free areas. To complete these surveys, an experimental study was carried out in quasi-natural conditions with engorged ticks placed in cages in the *garrigue*. It was shown that the nymphs moulted into adults within 3-4 weeks in July-August. However, they remain completely immobile in the litter, undergoing a behavioural diapause, until the following spring. On the other hand, almost all the nymphs released later, in September or October, died during the cold and rainy season. It was also observed that engorged females were able to survive the cold conditions up to 6-8 months before laying eggs. The study also suggested that there were predators of *H. marginatum*, likely ants and spiders, in the study site. The hosts monitored over all these years were not only infested by *H. marginatum*. The phenology of other tick species parasitizing horses (*Dermacentor marginatus, Haemaphysalis punctata, Rhipicephalus bursa*) and birds (*Ixodes frontalis, I. ricinus, H. punctata*) was also highlighted for this study site.

## Introduction

The two-host tick *Hyalomma marginatum* is one of the main vectors of Crimean-Congo Haemorrhagic Fever virus (CCFHv) (Bakheit et al., 2012). Larvae and nymphs infest mainly lagomorphs and birds, whereas adult ticks are known to parasitize domestic and wild ungulates (Morel, 2003; Walker et al., 2003). This tick species is established in northern Africa, Asia and southern Europe, from Portugal and Spain to Turkey (Santos-Silva and Vatansever, 2017). In Corsica, it has been present for at least 60 years (Morel, 1959) and its wide distribution on the island has been confirmed (Grech-Angelini et al., 2016). Until recently, there was no evidence of *H. marginatum* establishment in mainland France (Vial et al., 2016). However, during the past few years, such populations have been identified in at least seven French departments characterized by a Mediterranean climate. Between 2016 and 2022, thousands of *H. marginatum* adults have been collected, mainly on horses and cows (Bah et al., 2022; Bernard et al., 2024).

*Hyalomma marginatum* spreads from already infested areas to other favourable, yet uninfested, regions through the movement of infested hosts. These movements can involve horses (horse shows, trade, breeding, long distance horse riding, transfer from one to another livery stable etc.), cattle (movement between pastures or farms, trade), sheep (summer transhumance, trade) or wildlife such as wild boars – all of which infested by adult ticks. Larvae and nymphs are frequently observed on migrating birds captured in spring in various European countries, whether it be in the south, north or east of the continent (Jaenson et al., 1994; Jameson et al., 2012; Lindeborg et al., 2012; Capek et al., 2014; Nowak-Chmura, 2014; Vial et al., 2016; De Liberato et al., 2018; Pascucci et al., 2019; Portillo et al., 2021; Sormunen et al., 2021). This attests that larvae are active, at least in some of the infested areas, while birds are migrating towards Northern Europe. Birds therefore disseminate engorged nymphs, for which some will be able to moult into adults. Such adults have been recorded in many European countries where no established populations of *H. marginatum* have been observed, like Germany, Luxembourg, Netherlands, or Sweden (Chitimia-Dobler et al., 2019; Grandi et al., 2020; Weigand et al., 2020; Uiterwijk et al., 2021).

The establishment of *H. marginatum* in mainland France and the recent identification of CCHFv in some of these ticks collected close to the Spanish border (Bernard et al., 2024) suggest that the emergence of CCHF in the country may only be a matter of time. Preventing and controlling the disease first requires monitoring populations of the main tick vector. Therefore, it is essential to identify the temporal dynamics of *H. marginatum* to determine when the ticks are active and, ultimately, to assess their spreading potential. The purpose of this article is to describe three studies, carried out in the same study site, that led to (i) the determination of the seasonal variation of horse infestation by adult ticks, (ii) the collection of first data relating to seasonal variations of bird infestation by immature ticks, and (iii) the identification of the moulting kinetics of nymphs in quasi-natural conditions. The results provide the first insights into the lifecycle of *H. marginatum* in southern France which should be completed by observations made on different host species and at other sites within the henceforth extensive French distribution area of this tick species.

## Material and methods

### Study site and animals

In April 2016, a horse herd highly infested by adults of *H. marginatum* was identified 25 km north of Montpellier (Vial et al., 2016). This herd was chosen to study the seasonal variations of adult tick infestation. The monitored animals grazed freely 24 hours a day on a 200-ha clay-limestone land on *garrigue* (scrubland), Mediterranean forest and formerly cultivated fields now used as pastures, situated at an altitude of 330-350 m. During the 6-year-period of the survey, the herd size varied from 10 to 14 horses depending on births, deaths, introductions, departures or the temporary movement of mares for reproduction. These variations had to be considered for data analysis.

Leporidae (hares and rabbits), Erinaceidae (hedgehogs) and ground-dwelling birds are the hosts of *H. marginatum* immature ticks (Walker et al., 2003; Apanaskevich and Horak, 2008; Santos-Silva and Vatansever, 2017). Hares have rarely been observed on the land, and no rabbit warren was located during the survey. To assess the phenology of immature stages, it was therefore decided to examine birds.

The development and survival of *H. marginatum* during the winter period were also studied on this land. Engorged nymphs and engorged females were placed in cages protected by an enclosure set up on one of the property’s non-wooded areas (*garrigue*). These ticks were obtained from breeding established in 2018 at the Baillarguet^1^ insectarium from engorged females collected on horses within the monitored herd. The adult ticks were fed on goats and the immatures were fed on rabbits in accordance with the procedures described in the ministerial authorisation for “feeding ticks on vertebrates for laboratory rearing” n° APAFIS#6304-2016080315594914 v4. Engorged ticks were kept in climatic chambers at 10°C or 12°C and 85 % humidity to prevent them from developing in the laboratory before being released in the field.

The study site has a typical Mediterranean climate, characterized by warm and dry summers, mild winters, and rainfall concentrated over a few days in autumn (principally) and spring. Meteorological data are recorded by Météo-France at a place (altitude: 214 m) situated 13 km away from the study site^2^ (Supplementary Data S1). During the study period, the average annual temperature was 14.5°C, the maximum daily temperature ranged 27.6°C to 29.5°C, with the highest observed temperature (41.8°C) recorded in 2019. The annual rainfall was highly variable, from 663 to 1438 mm. Rainfall exceeding 1 mm was recorded for an average of 72 days a year, and 80% of annual rainfall was recorded in fewer than 30 days. The entire area, from this place to the study site, is heavily infested by *H. marginatum*.

### Study periods

Tick collection on horses was performed from April 2016 to April 2022. Horse examination was usually carried out every 6-14 days during the *H. marginatum* infestation peak in spring and summer (Grech-Angelini et al., 2016), and every 15-30 days during the rest of the year. The survey was interrupted on different occasions: between May 2017 and April 2018 (funding interruption), from February to mid-August in 2020 (suspension of field activities during the Covid-19 pandemic), from May to September in 2021 (treatment of the animals with different experimental acaricide preparations). Overall, tick infestation on horses was monitored for about 50 months over the six years, with an emphasis in March-April, at the onset of infestation.

Birds were captured and examined in 2016 (from the end of June to the end of October), in 2017 (July and August) and in 2019 (from mid-March to mid-July). Nets were installed for 2-3 consecutive days approximately every 3-4 weeks, except when rain prevented bird captures that were therefore cancelled.

Finally, the development of engorged nymphs and females was studied in quasi-natural conditions between July 2019 and June 2020.

### Tick collection on horses and birds

All ticks attached to the examined horses were collected and identified back in the laboratory. Some of them were kept alive until identification and then deep-frozen at -80°C for further analysis (Bernard et al., 2024; Joly-Kukla et al., 2024). On other occasions, ticks were stored in 70% ethanol and discarded after identification.

It was impossible to maintain a constant interval between two successive tick counts as the horses could not always be found on the scheduled examination date. Moreover, the number of examined animals constantly varied. All these variations had to be considered for data analysis. In order to homogenize the results and consider these constraints, it was decided to calculate the daily number of *H. marginatum* ticks attached per horse between two successive tick collections (DNT). The total number of *H. marginatum* collected during a tick collection was divided by the number of animals examined and by the number of days separating this collection from the previous one.

Birds were caught in the study site using 210 m of nets (5 sets of 25-70 m). These were installed in narrow forest paths and opened about one hour before sunrise for 4-5 hours, being checked every 30 minutes. Captured birds were identified, ringed, weighed, examined for tick infestation and then released. All attached ticks were collected and kept alive until identification in the laboratory. Bird capture was authorised by the “Centre de recherche sur la biologie des populations d’oiseaux (CRBPO)” (Research Centre on Bird Populations Biology) and the Muséum national d’Histoire naturelle of Paris, and carried out by an experienced ornithologist and bird bander.

Ticks were identified by experienced acarologists using the keys and descriptions of Walker et al. (2003) and Pérez-Eid (2007). Two methods were used to confirm the identification of the *H. marginatum* immatures. In 2016, the fully engorged nymphs were kept in a climatic chamber (25°C and 85% RH) until metamorphosis which allowed the morphological identification of the moulted adults. In 2017, part of the collected *H. marginatum* larvae and nymphs were analysed and determined in a phylogeographic study. DNA extraction, amplification and sequencing of two mitochondrial markers (12S rRNA and Cytochrome Oxidase 1) were carried out and are described in Giupponi et al. (2025).

### Development of engorged ticks in quasi-natural conditions

The cages placed in the field with engorged ticks were the same as those previously used in Burkina Faso and Madagascar for similar experiments with the hard tick *Amblyomma variegatum* (Stachurski et al., 2010; Rahajarison et al., 2014): they are 50cm-side metal frames cubes, covered with mosquito nets on all sides except the bottom. The lower frame was buried 3-5 cm into the ground to prevent ticks from escaping. The cages were placed on shrubs like thymus, rosemary, *Genista scorpius* or large tufts of grass and contained natural plant litter to ensure that the ticks could find micro-habitats favourable to their survival and development.

Four cohorts of 4 or 6 cages, each containing 26 to 30 engorged nymphs and one or two engorged females (according to the number of ticks available at the time of installation), were studied. The four cohorts were respectively installed in the field on 10/07, 14/08, 18/09 and 24/10 of 2019. The first three dates correspond to the period during which the hosts were infested by *H. marginatum* larvae and nymphs, as observed during the bird survey. To verify whether the tick development (nymphs moulting and egg-laying) occurs rapidly, the first two cages were examined 3 and 6 weeks after the release of the engorged ticks for the 4 cohorts. A third cage was checked 9 weeks (for cohort 3) or 12 weeks (for cohort 4) after the installation of the cages because nymph moulting had not yet started 6 weeks after release for these two cohorts. The last cages were examined in the next spring (end of March and mid-June for all cohorts, and end of April for cohorts 3 and 4) to verify whether the unfed adults, moulted before winter, were still alive, and whether the unmoulted nymphs had been able to moult with the increase of the temperatures. The cages were examined destructively: plants were removed and the ticks were searched for, first at the base of the shrubs or grass tufts, then in the litter, and finally in the first centimetres of soil, once the plant debris had been removed. The soil removed from the cage was carefully examined. Finally, the cage itself was removed, and the ticks were searched for in the hem of the mosquito nets covering it. The collected ticks were brought back to the insectarium where the unmoulted nymphs and females which had not started to lay eggs were maintained in a climatic chamber (25°C, 85% RH) to verify whether they were alive and still able to develop or lay eggs.

## Results

### Horse infestation

More than 13,800 ticks were collected from the horses during the study period (Table 1). *Hyalomma marginatum* was the most abundant species (85% of the collected ticks), mostly attached to the perineal area, inguinal region, udder and sheath. *Rhipicephalus bursa* (6% of the collected ticks) were collected from the same anatomical regions. Adult *R. bursa* parasitized the horses from March to August, with a peak in June-July, while nymphs were collected from December to April. The number of collected *R. bursa* decreased drastically between 2016 and 2021. *Dermacentor marginatus* (8%) and *Haemaphysalis punctata* (1%) were found during the cool season, between September-October and April-May (Supplementary Material S2). The former species was found almost exclusively on the horses’ mane whereas the latter was mainly attached to the chest. Finally, a few *Rhipicephalus* of the *sanguineus* group and two *Ixodes ricinus* females were also found, mainly during the first years of monitoring.

**Table 1.** Number and species of the ticks collected from the monitored horses during the various study periods.

| Year | Period | *Hyalomma marginatum* |  | *Dermacentor marginatus* |  | *Haemaphysalis punctata* |  |  | *Rhipicephalus bursa* |  |  | *Rhipicephalus* of the *sanguineus* group |  | *Ixodes ricinus* |  |
| --- | --- | --- | --- | --- | --- | --- | --- | --- | --- | --- | --- | --- | --- | --- | --- |
|  |  | M | F | M | F | M | F | N | M | F | N | M | F | M | F |
| 2016 | April-Dec | 957 | 910 | 92 | 73 | 14 | 15 | 0 | 188 | 254 | 25 | 6 | 6 | 0 | 1 |
| 2017 | Jan-April | 475 | 295 | 199 | 207 | 8 | 39 | 0 | 29 | 37 | 37 | 5 | 2 | 0 | 1 |
| 2018 | April-Nov | 1427 | 1609 | 13 | 9 | 2 | 8 | 0 | 71 | 111 | 2 | 0 | 1 | 0 | 0 |
| 2019 | Jan-Dec | 1629 | 1727 | 99 | 142 | 5 | 33 | 1 | 31 | 50 | 6 | 0 | 0 | 0 | 0 |
| 2020 | Jan-Feb; Aug-Dec | 23 | 37 | 57 | 86 | 2 | 9 | 0 | 0 | 0 | 0 | 0 | 0 | 0 | 0 |
| 2021 | Jan-May; Sep-Dec | 1180 | 1015 | 72 | 78 | 4 | 14 | 0 | 2 | 4 | 0 | 2 | 0 | 0 | 0 |
| 2022 | Jan-april | 265 | 145 | 10 | 17 | 3 | 7 | 0 | 1 | 0 | 0 | 0 | 0 | 0 | 0 |

Focusing on *H. marginatum*, few ticks were collected in January and February. Adults of this species infested horses mainly between March and the end of August (Figure 1). Generally, male ticks started to attach a slightly earlier than females, but the latter were more numerous during the second part of the infestation period (Supplementary S2; Data D1). During the first seven days of March, an average of 0.18 ticks attached daily to each horse (DNT). This pattern was almost similar over the course of the monitored years: 0.16 in 2017, 0.24 in 2019, 0.14 in 2021, 0.18 in 2022.

**Figure 1.**
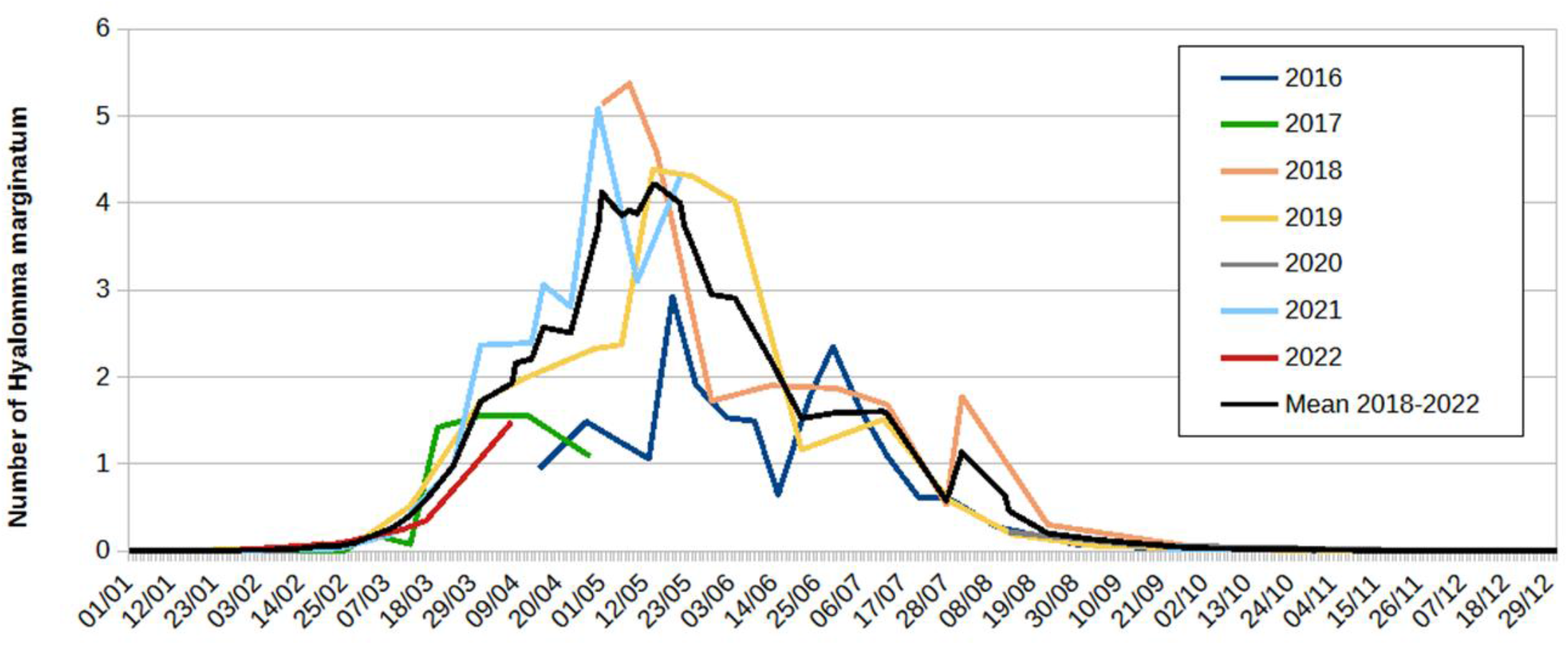
Number of *Hyalomma marginatum* adults attached daily to the monitored horses during the various study periods. The values were calculated as follows: the number of *H. marginatum* collected during one tick collection session was divided by the number of animals examined and by the number of days separating this collection from the previous one.

As the infestation observed in the first survey year (April 2016 to April 2017) appeared lower than those recorded in subsequent years before levelling off, the mean DNT (black curve of Figure 1) was calculated with the data of the last five years (2018-2022). This mean DNT reached 0.5 ticks on 15^th^ March, 1.0 on 25^th^ March and 2.0 on 9^th^ April. It remained higher than 2.0 until 15^th^ June and higher than 1.0 until 20^th^ July. The mean DNT peak (4.0 ticks attached daily) was observed during the first three weeks of May. This daily infestation represented a tick burden of approximately 330 *H. marginatum* per year and per horse, whereas the infestation reached roughly 200 ticks per animal during the first year of monitoring (April 2016 to April 2017). During the three years in which the horses were regularly examined throughout summer (2016, 2018 and 2019), an additional minor increase in infestation was observed in July or August after an initial reduction of the DNT.

### Bird infestation

Birds were captured during 30 non-consecutive days: 15 in 2016, 6 in 2017 and 9 in 2019.

The data were aggregated per week over a fictional year (Figure 2). A total of 317 birds (1-25 per day of capture) of 25 different species were examined (Table 2). Those most frequently found in the nets were the European robin, the common blackbird, the Eurasian blackcap and the great tit. These four species were observed on site throughout the year. Others (flycatcher, nightingale) were only captured from mid-May to mid-September. Several species were caught only once or twice (song thrush, Eurasian wren).

**Table 2.** Species and number of birds examined and infested by ticks of all species during the study.

| Name of the bird | Number |  | Name of the bird | Number |  | Name of the bird | Number |  |
| --- | --- | --- | --- | --- | --- | --- | --- | --- |
|  | examined | infested |  | examined | infested |  | examined | infested |
| European robin<br>(*Erithacus rubecula*) | 81 | 15 | Eurasian blackcap<br>(*Sylvia atricapilla*) | 31 | 0 | Cirl bunting<br>(*Emberiza cirlus*) | 2 | 0 |
| Common blackbird<br>(*Turdus merula*) | 56 | 29 | Common firecrest<br>(*Regulus ignicapilla*) | 19 | 0 | Rock sparrow<br>(*Petronia petronia*) | 2 | 0 |
| Great tit<br>(*Parus major*) | 23 | 3 | Long-tailed tit<br>(*Aegithalos caudatus*) | 13 | 0 | Garden warbler<br>(*Sylvia borin*) | 2 | 0 |
| Common nightingale<br>(*Luscinia megarhynchos*) | 21 | 4 | Sardinian warbler<br>(*Curruca melanocephala*) | 10 | 0 | Eurasian jay<br>(*Garrulus glandarius*) | 1 | 0 |
| Western subalpine warbler<br>(*Curruca iberiae*) | 12 | 1 | Short-toed tree creeper<br>(*Certhia brachydactyla*) | 9 | 0 | Melodious warbler<br>(*Hippolais polyglotta*) | 1 | 0 |
| Eurasian chaffinch<br>(*Fringilla coelebs*) | 11 | 2 | European pied flycatcher<br>(*Ficedula hypoleuca*) | 6 | 0 | Eurasian blue tit<br>(*Cyanistes caeruleus*) | 1 | 0 |
| Dunnock<br>(*Prunella modularis*) | 3 | 1 | Common chiffchaff<br>(*Phylloscopus collybita*) | 4 | 0 | Western Orphean warbler<br>(*Curruca hortensis*) | 1 | 0 |
| Song thrush<br>(*Turdus philomelos*) | 1 | 1 | Great spotted woodpecker<br>(*Dendrocopos major*) | 4 | 0 | Common redstart<br>(*Phoenicurus phoenicurus*) | 1 | 0 |
|  |  |  | Eurasian wren<br>(*Troglodytes troglodytes*) | 2 | 0 |  |  |  |

**Figure 2.**
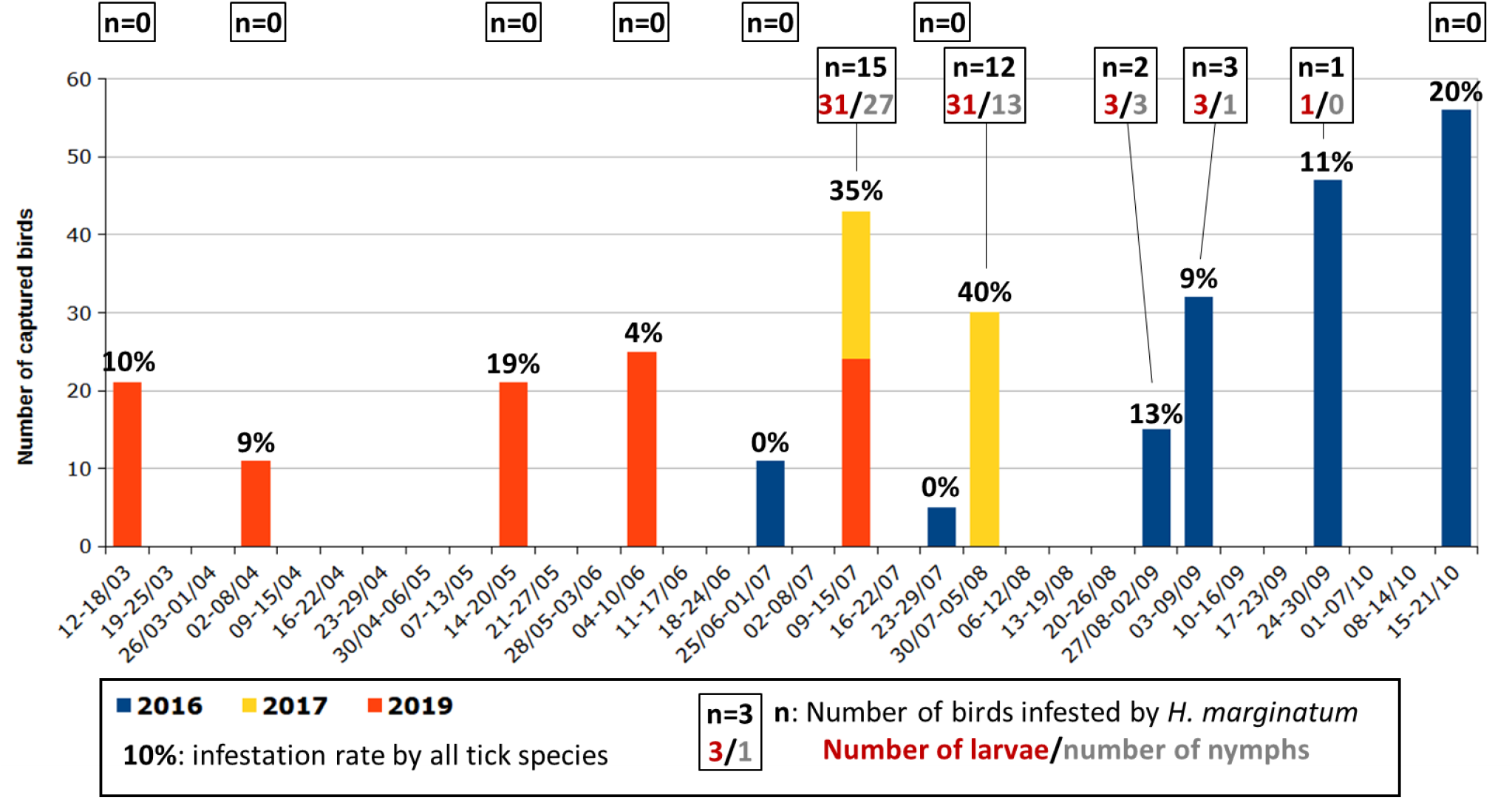
Number of birds examined during the surveys and tick infestation rate of these birds with special emphasis on *Hyalomma marginatum* immature infestation.

Ticks were found on 56 birds of eight species (Table 2). The overall infestation rate was 18%, and varied over the year (0-40%). In total, 167 ticks of four species were identified, as well as two Ixodidae larvae too damaged to be identifiable (Supplementary Material S3). The most abundant tick species was *H. marginatum* with 44 nymphs (NN) and 69 larvae (LL), collected from 32 birds of seven different species (all the infested bird species except the Dunnock). *Ixodes frontalis, I. ricinus* and *H. punctata* were also collected. There were generally up to five ticks per bird (mean: 3; median: 2). The only exceptions were *H. punctata* larvae (14 were found on a blackbird), and *H. marginatum* immatures with six birds (five blackbirds and one robin) infested by six to 18 ticks. Blackbird was the most infested bird species, harbouring 83 of the 113 *H. marginatum* (73%), as well as 95% of the 19 *H. punctata* and 48% of the 25 *I. frontalis*. However, no *I. ricinus* was found on a blackbird. Three of the four *Hyalomma* nymphs collected in 2016 were engorged enough to moult into adults when kept in the climatic chamber. They were all *H. marginatum*. The mitochondrial markers of 28 of the *Hyalomma* immatures collected in 2017 (3 of the 48 larvae and 25 of the 39 nymphs) were analysed (Giupponi at al., 2025). They were all *H. marginatum*, confirming the morphological identification.

Co-infestations (*I. ricinus* and *I. frontalis* or *H. marginatum* and *H. punctata*) were occasionally observed. Each tick species displayed a particular phenology. *Haemaphysalis punctata* (2 NN and 17 LL) were collected mainly in July and August, except for one nymph collected in May. *Ixodes ricinus* (2 NN and 8 LL) were observed in September and October. Larvae of *I. frontalis* (n = 13) were only collected in October while nymphs of this species (n = 9) were collected in October and from March to June, as well as 3 females in September and May (Supplementary S3).

No immature *H. marginatum* were collected in May (21 birds examined) or in June (36 birds examined). They were collected from the beginning of July to the end of September, mainly during the first month (Figure 2). Larvae and nymphs of *H. marginatum* were often observed on the same birds. These immatures were first observed on 10^th^ July in 2019, on 11^th^ July in 2017 but only on 31^st^ August in 2016. That year, only five birds have been captured in July, and none of them were blackbirds, whereas 19 of the 32 birds (59%) found infested by *H. marginatum* during the study were blackbirds. More precisely, 21 of the 32 (66%) blackbirds examined from July to September were infested by *H. marginatum* immatures. This proportion reached 33% (2 of 6) for the Eurasian chaffinch, 27% (3 of 11) for the common nightingale, 22% (2 of 9) for the great tit, 13% (1 of 8) for the western subalpine warbler and 9% (4 of 45) for the European robin. The only song thrush caught was also infested by *H. marginatum*.

### Moulting of *H. marginatum* nymphs in quasi-natural conditions

The number of ticks found in the cages was highly variable from one cage to another, with a general (but not systematic) decrease over the time (Figure 3 and Data D2). Similarly, the fate of the engorged nymphs varied greatly from a cohort to another. Predated ticks (mainly nymphs, but also two unfed adults) were sometimes collected in the cages. In one cage of cohort 2 in particular, almost a third of the nymphs had been predated. In these cases, the cuticle was pierced and the tick was empty (nymph), or only part of the scutum and legs were found (adult) (Supplementary S4).

**Figure 3.**
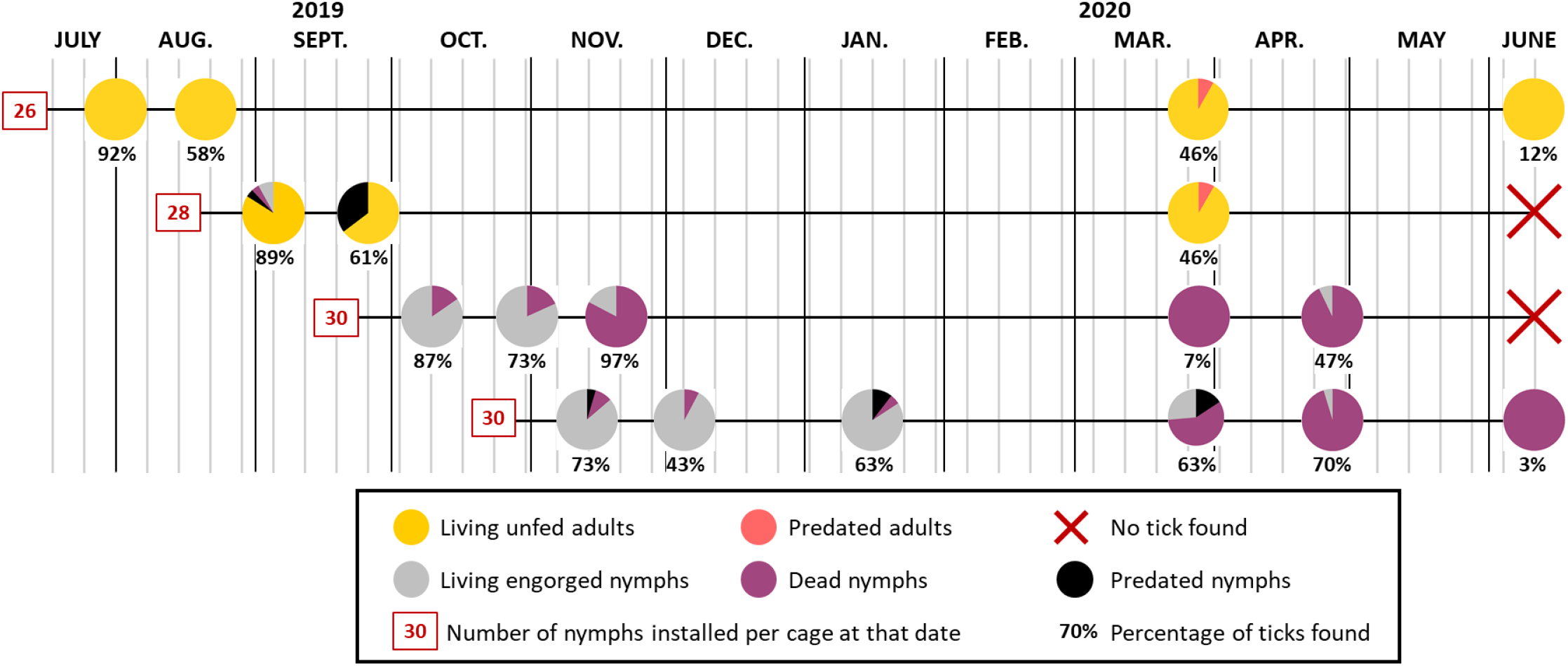
Graphic representation of the status of the ticks found in the cages where batches of 26 to 30 engorged nymphs, according to the cohort, had been placed. Some of the ticks were found killed by redators (ants, spiders, beetles): cuticle pierced, tick partly eaten. The not predated nymphs were placed at 25°C and 85% RH in a climatic chamber at the CIRAD insectarium to verify whether they were still able to moult, and therefore alive, or dead.

Three weeks after the engorged nymphs of cohort 1 had been placed in the field, all the ticks found in the first cage (24 of the 26 installed ticks) were adults. However, the metamorphosis was recent, as the legs of some ticks were still light-coloured and the pale rings at the outer ends of the segments were not yet visible. An additional three weeks later, all the ticks found were adults with clearly bicoloured legs. All the adults in these two cages were completely immobile when they were found.

Twenty-one of the 25 ticks (out of 28 installed) found in the first cage of cohort 2 were adults, while three had not yet completed their metamorphosis. Two of these three nymphs moulted in less than a day when placed in the insectarium, while the last one died. As in cohort 1, all living ticks found three more weeks later (many nymphs had been predated) were adults, but they remained completely inactive in the cages.

The following spring, at the end of March, unfed adult ticks moved into the cages as soon as the cages were checked. They climbed up the mosquito nets on the side where the human was. No living ticks were found immobile on the ground on which only predated adults were found. At this time, the adults were 200-240 days old. At the beginning of June, only a few active ticks (around 310 days old) were still found in the cage from cohort 1. The mosquito nets were badly damaged by that time, as a result of the adverse weather conditions, particularly sun exposure, which had been affecting them for almost a year.

The pattern was completely different for ticks belonging to both cohorts 3 and 4. All the ticks found in the cages were nymphs (no adult *H. marginatum* was found in the cages of these two cohorts). In the first four cages of cohort 4, some of the nymphs were still mobile when found. Placed in a climatic chamber, only few nymphs of these two cohorts were able to moult. The median duration of the metamorphosis was 10, 6 and 5 days, respectively, for the nymphs in the first three cages of cohort 3, and 18-21 days for those in the first four cages of cohort 4.

A large proportion (83%) of the nymphs of cohort 3 collected on 19/11 were dead while only 6-10% of the nymphs in cohort 4, collected on 13/11, 04/12 and 15/01, were dead. However, the mortality rate of the nymphs from these two cohorts was high the following spring: 87% of those collected between March and June were dead or unable to develop when put in the climatic chamber. In April, only one of the 14 nymphs from cohort 3 succeeded to moult within 24 hours when put at 25°C and 85% RH. This was also the case for only one of the 21 nymphs of cohort 4 collected in April. Metamorphosis occurred in 8 days. It took 2 or 4 weeks for these 2 ticks to emerge from their exuvia.

The ticks of these two cohorts often appeared to be in good condition when placed in the climatic chamber, but many were unable to emerge from the nymph’s cuticle, even if they had completed their metamorphosis. This exuvia was much thicker than that observed during normal metamorphosis. When they were voluntarily removed from the exuviae after 20-30 days, it was observed that the adults were poorly formed. In particular, the legs were twisted, the articles shorter than normal and the joints were deformed (Supplementary S5). As for the few adults that were able to emerge from the exuvia in these cohorts, nearly all died within a few days, without ever showing any ability to respond to stimuli (breathing, light).

### Egg-laying by *H. marginatum* females in quasi-natural conditions

Only one of the 12 engorged females placed in the cages of cohorts 1 and 2 was found (Figure 4 and Data D3). It had begun to lay eggs. However, it was not in the litter nor on the ground but in the hem of the mosquito net. The situation was completely different for females belonging to the cohort 4: all were found, and only one, being 155 days old, was dead (in March). Those found in November-January needed 6-17 days in the climatic chamber to start laying eggs, whereas those found in March-April needed 1-2 days. The two females observed in the cage in June (230 days old) had started laying eggs in the field. The situation for cohort 3 was intermediate: all the females were found in October-November and some had started to lay eggs, while others needed 2-11 days in climatic chamber to start oviposition. Only two females (one predated and one alive, which quickly started to lay eggs in the climatic chamber) were found in spring.

**Figure 4.**
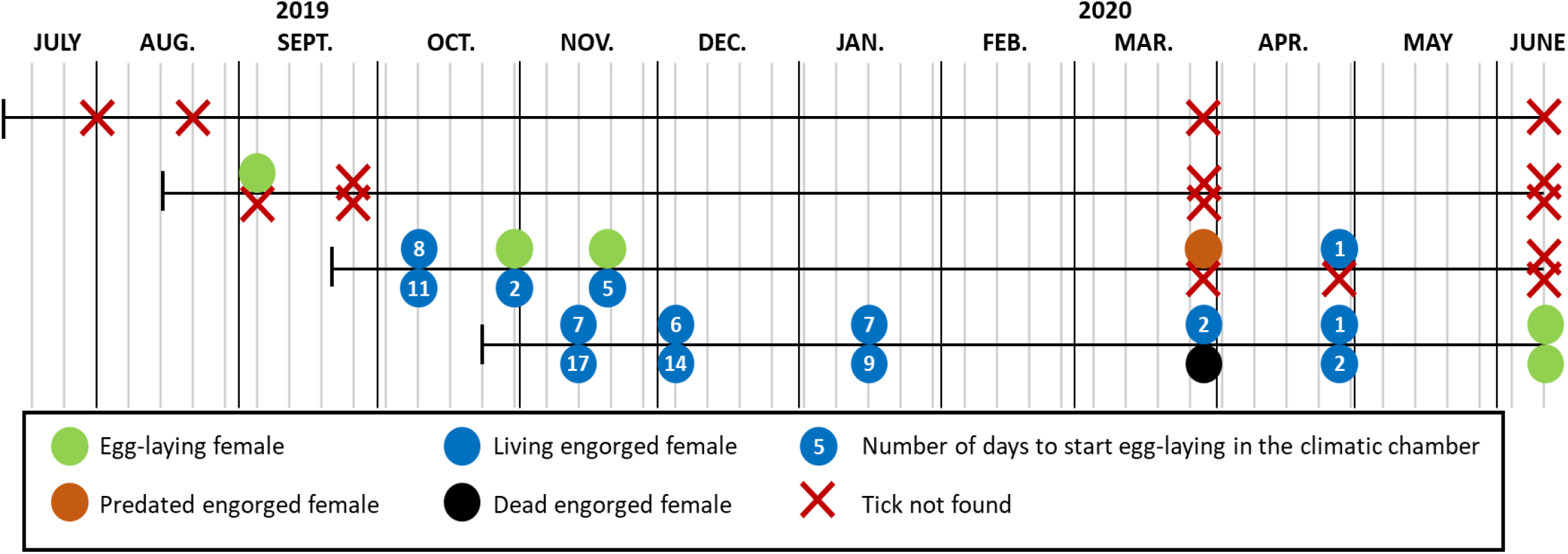
Graphic representation of the status of the engorged females from the 4 cohorts found in the cages where they had been placed. The females which had not started to lay eggs were placed at 25°C and 85% RH in a climatic chamber at the CIRAD insectarium to verify whether they were still alive and able to lay eggs. One female was killed (cuticle pierced) by a predator, probably a spider.

## Discussion

Before discussing the main results obtained from this survey on *H. marginatum* in this site of southern France, it should be mentioned that some limitations were identified in the study. For different reasons, the ticks collected from the monitored horses do not represent all those that infested them during the study period. First, the horses selected for this study were only minimally handled because of their semi-wild lifestyle. Consequently, some individuals were sometimes reluctant to undergo a thorough examination and tick collection. Secondly, the interval between two tick samplings sometimes exceeded the duration of the female tick’s blood meal. Some of those females were thus able to attach, engorge and drop off between two tick collections. Thirdly, from the end of 2019, when tick collection was occasionally suspended for more than 3-4 weeks and infestation became high, the owner treated the most infested animals with an acaricide, preventing new ticks from attaching to these horses. They were therefore removed from the survey until new ticks attached to them. The acaricide was on the other hand occasionally applied, during the cool periods, to the manes of the highly parasitized horses to prevent wounds and extensive scabs due to high infestation by *D. marginatus*. Finally, as the objective of the study was to ascertain the seasonal variations of *H. marginatum* infestation, tick sampling was carried out with reduced frequency during the cool season, between September and February. Consequently, even if these results do not represent the core of this study, the number of active ticks in autumn and winter, mainly *D. marginatus* and *H. punctata*, was probably underestimated compared to their actual proportion in the tick population present in the site.

It appeared that *H. marginatum* was the most abundant tick on the monitored horses. Its abundance seemed to increase during the first two years of the survey, before stabilising from 2018 onwards. This rapid increase suggests that the establishment of *H. marginatum* on the farm was relatively recent. Over the same period, the number of *R. bursa* collected on horses dramatically decreased. The suspension of tick collections during the summers 2020 and 2021, because of Covid-19 pandemic or acaricide trials, may have contributed to the apparent decrease in *R. bursa* infestation since the infestation peak by this species is observed at that period. It is also remarkable that very few *Rhipicephalus* of the *sanguineus* group were collected from the horses whereas 34 of them were found on a single wild boar that was shot by a hunter on the land in the summer of 2016. This wild boar was also infested by six *H. marginatum* (5 males and 1 female) whereas 1.5 *H. marginatum* attached daily to each horse during this period. This is an indication that these two tick species probably do not have the same predilection host.

According to the herd owner, *H. marginatum* was observed on the horses 3-5 years before the beginning of the survey. It has not been possible to determine how the tick was introduced in the farm. No horse was purchased in the years prior to the start of the survey, but a few mares have been sent to other farms each year for 2-3 months in spring for breeding. They may have come back infested with *H. marginatum* from one of these farms. Migratory birds (nightingale, song thrush) coming from Spain or North Africa nest on the study site every year. Due to the very small number of migratory birds caught in spring (no nightingale was examined before mid-May, for example), it is not possible to know whether the introduction of *H. marginatum* nymphs was likely to occur through this route. When discussing the spread of *H. marginatum* in Eastern Europe, Gray et al. (2009) wrote that “it is debatable (…) because very small numbers of ticks, all immature stages, would be involved.” They rather consider that “establishment of *H. marginatum* (*would*) mainly result from the introduction of adult females”.

Infestation of horses by *H. marginatum* adults started slowly at the end of February or at the beginning of March. The number of ticks which attached daily increased until the beginning of May to reach roughly four ticks per host per day, *i*.*e*., about 30 *H. marginatum* per week in the final years. From June, the number of active adult ticks in the environment and infesting animals decreased, dropping back to nearly zero by early September. This temporal pattern was repeatedly observed over the six years, suggesting predictable temporal variations in *H. marginatum* dynamics. In 2016, 2018 and 2019, a new infestation peak was observed in late summer (Figure 1), when the animals were free to graze on a pasture that had been deferred a few months earlier to preserve the grass resource. This points out that adult ticks, which could not previously attach, were still alive and active in the late summer. In autumn and winter, a few *H. marginatum* adults, mainly males, had been found on the hosts, but this was rather unusual. Only 0.5% of the 11,694 *H. marginatum* collected from the horses during the study were found between mid-September and mid-February.

*Hyalomma marginatum* males have never been reported to produce attraction-aggregation-attachment pheromone similar to that emitted by *Amblyomma hebraeum* males (Rechav et al., 1977) or *A. variegatum* males (Norval and Rechav, 1979) after 3-4 days of attachment. The fact that the males attach slightly earlier than the females is more likely due to differences in the level of stimuli that trigger host-seeking activity at the end of winter. The data collected did not permit determination of the main stimulus (daylength, temperature exceeding a given threshold, another phenomenon) inducing the host-seeking activity of *H. marginatum*.

During the present study, *H. marginatum* nymphs were observed on birds from 10^th^ July. As this species is a two-host tick whose immature stages infest their host for about 3 weeks (Gargili et al., 2013), it is assumed that larvae were active from the second half of June. However, by the end of August, the number of *H. marginatum* found on the birds dropped drastically, suggesting that there were no longer active/living larvae in the environment. The infestation period by *H. marginatum* larvae on this site lasts thus about 3 months (mid-June to mid-September), shorter than the period estimated by Estrada-Peña et al. (2011) for whom “larval recruitment may be feasible from March to September”. Larvae were therefore not yet active between March and May, when the migratory birds arrive from Africa or Spain. If this phenology is also observed in all French regions where the tick is established, which has to be studied, birds parasitised by *H. marginatum* immatures flying to northern Europe could not have been infested during their passage through southern France. They were thus infested by larvae further south, for example in Morocco, where birds were found to be infested by *H. marginatum* immatures as early as April (Palomar et al., 2013). To the best of our knowledge, the period of activity of *H. marginatum* larvae in Spain has not been identified. It may also be different between the southern regions of the country (Andalusia, Extremadura, Castilla–La Mancha) and those close to the south France border, such as Catalonia.

Birds infested by *H. marginatum* were mainly ground-dwelling birds, spending a large part of their life on soil: blackbird and probably song thrush (although only one was examined), nightingale, robin or, to a lesser extent, chaffinch. Other ground-dwelling birds, such as partridge and quail, which were not caught in this study, are also likely to be good hosts for *H. marginatum* larvae (Hoogstraal et al., 1963; Hosseini-Chegeni et al., 2019). As roughly two-thirds of the blackbirds were infested by *H. marginatum* immatures in summer, it is suggested that they could be a good sentinel for determining whether the species is established in a region and whether larvae are active in the area. They are generally abundant and widespread in the suitable habitats, and they are heavily parasitised by this tick species.

While the core information of this study concerns *H. marginatum*, it is interesting to note that none of the blackbirds examined in September and October, when the migratory birds arrive in the Mediterranean region from the north, were infested by *I. ricinus* larvae or nymphs. It suggests that the vast majority of those captured at that time were birds that had grown up in the area. Conversely, the number of robins caught increased dramatically from the end of September (28 in 27-29/09 and 26 in 19-21/10 instead of 1-3 per day at the other periods). Six of them (11%) were infested by *I. ricinus* larvae or nymphs. However, only two *I. ricinus* females were found on the monitored horses (one in April 2016 and another in March 2017). This indicates that despite the autumnal introduction of *I. ricinus* larvae and nymphs, the species does not seem to be able to establish in this site, where the climate is most likely unsuitable. Moreover, during surveys carried out between 2017 and 2022 to determine the distribution of *H. marginatum* (Bah et al., 2022), it was found that the two species rarely coexisted: *H. marginatum* was found in hot, dry areas, and *I. ricinus* in wetter, temperate sites. In Corsica, *I. ricinus* was hardly found, uniquely at altitudes of over 600 m in the coolest areas (Grech-Angelini et al., 2016).

The infestation rate of blackbirds (52%) or robins (19%), as well as the overall infestation rate of the 317 birds captured on the site (18%) was much higher than the rates generally observed in studies focusing on migratory birds. Hoogstraal et al. (1963) observed ticks on 3% of the birds captured in 1959-1961 “in Egypt while en route from Asia and eastern Europe to tropical Africa”. Pascucci et al. (2019) found that 7% of the 3444 birds examined during spring migration in Ventotene island, in the central Tyrrhenian Sea, were infested by ticks. De Liberato et al. (2018) found ticks on 0.3% of the birds examined in March-May 2013 and 2014 in various western coastal sites of central Italy. Bacak et al. (2023) found ticks on 6% of the 10,650 birds examined in the Turkish Thrace during the migrating periods of 2020-2022. This was very probably because almost 60% of the birds examined in the study presented here were ground-dwelling birds, which was generally not the case.

Except for the cages examined in June and one cage from cohort 3, 43-97% (mean: 67%) of the ticks placed in the cages were found up to eight months after installation in the field. The disappearance of some ticks may be due to an imperfect examination of the litter, especially when the ground was muddy and sticky, just after a storm. This may also be due to ticks escaping through the holes in the mosquito netting, especially during the final months of the study, when the unfed adults were very active and the nets were badly damaged from sun exposure.

However, some ticks had been predated. The most often mentioned predators, frequently found in the cages, were ants. In certain regions of Australia favourable to *Rhipicephalus microplus*, the absence of the tick has been attributed to the predatory action of ants (Wilkinson, 1970). Ants of the genus *Solenopsis* have also been observed feeding on engorged female *R. (Boophilus)* or *Amblyomma* (Burns and Melancon, 1977; Butler et al., 1979; Petney et al., 1987; Barré et al., 1991). More recently, it was shown that the number of questing ticks (mainly *I. ricinus*) was negatively correlated with the size of red wood ant nests (Zingg et al., 2018). In any case, when ants are involved, they generally do not leave the corpse of the predated ticks in the cage but take it to their anthill (Mwangi et al., 1991). This could explain why some of the ticks had disappeared from the cages. To date, no data has been reported on tick-predating ant species in France. As for the ticks found pierced and empty, they were probably preyed by other predators such as spiders or beetles (Samish and Alekseev, 2001; Vinogradov et al., 2023), which were also found in the cages. The predatory activity is extremely variable and unpredictable in the field (Wilkinson, 1970; Petney et al., 1987; Stachurski et al., 2010). This was also observed in this study. There are thus predators of *H. marginatum* in the environment, apparently more active in hot weather, but their importance in terms of reducing tick populations is unknown.

The cage experiment showed that, in summer, *H. marginatum* nymphs moult very quickly into adults, which remained inactive. This indicates that the adult ticks undergo a behavioural diapause, already described for this species by Belozerov (1982). This has also been observed for *Amblyomma variegatum* adults in Burkina Faso (Stachurski et al., 2010). In this latter case, the end of the diapause was observed with the onset of the rainy season. The situation with *H. marginatum* is likely to be different as there is no significant increase in rainfall in early March, when the adults become active. Behavioural diapause is characterised by the suppression of host-seeking activity in unfed ticks. For Belozerov (1982), this is a “pre-adaptive behaviour which precedes the actual onset of unfavourable environmental conditions”, mainly regulated by the photoperiod. It enables the synchronisation of the life-cycle with favourable seasons of the year. The host-seeking activity of unfed *H. marginatum* adults in March would thus be triggered by the increase in daylight hours, which announces the rise in temperature, which in turn favours eggs incubation.

At 25°C, engorged *H. marginatum* nymphs moult in 18.6 days (median time; unpublished data). Nymphs from cohort 4 required 18-21 days, showing virtually no development while in the field. Some were even still moving despite spending up to 155 days in the cages. In contrast, cohort 3 nymphs moulted within 5-10 days at 25°C, indicating that their metamorphosis had already begun in the field. However, many nymphs in cage 3 of this cohort were found dead mid-November, unlike those of cohort 4 collected at the same time. Most cohort 4 nymphs remained even alive during the coldest winter period. An environmental factor likely disrupted cohort 3 nymphs between late October and mid-November. Heavy autumn rainfall is suspected to have hindered successful moulting of these nymphs which, when the development had begun, are completely immobile and vulnerable to submersion. Conversely, active nymphs can escape unfavourable conditions. This may explain the higher survival of cohort 4 nymphs compared to cohort 3 ticks. The survival of cohort 4 nymphs declined however sharply later, to reach 30% in March and 5% in April. Overall, adverse weather conditions appeared to severely disrupt metamorphosis, causing death or deformities in emerging adults.

*Hyalomma marginatum* unfed adults can survive winter because of behavioural diapause, unlike engorged nymphs. In contrast, *A. variegatum* unfed adults survive the cool season in Burkina Faso through behavioural diapause (Stachurski et al., 2010), whereas, in the Malagasy highlands, engorged nymphs are able to survive the very cold season, resuming their development up to 20 weeks (Rahajarison et al., 2014). In both cases, the ticks survived during the dry season, suggesting that rainfall, or a combination of rainfall and low temperature, may prevent survival of engorged *H. marginatum* nymphs in continental France. This hypothesis requires experimental confirmation.

The experiment carried out with cages showed that engorged *H. marginatum* females can survive the unfavourable conditions, unlike engorged nymphs. Some of the females placed in the cages in autumn were able to lay eggs up to 230 days later. Egg-laying was likely delayed by low temperatures rather than due to true diapause. It is unclear whether the eggs laid in autumn can survive winter and hatch in the following spring, joining those of the next generation. However, this may not reflect natural conditions as very few females feed after early September.

These various studies have made it possible to identify the cycle of *H. marginatum* in this site of southern France (Figure 5). Adult ticks emerge from their behavioural diapause from late February or early March and then infest their ungulate hosts. In the study site, these hosts are mainly horses, but the tick can also parasitise wild boars and possibly the few roe deer present. In the vicinity of the study site, there are also sheep flocks and cattle herds that are infested by *H. marginatum*.

**Figure 5.**
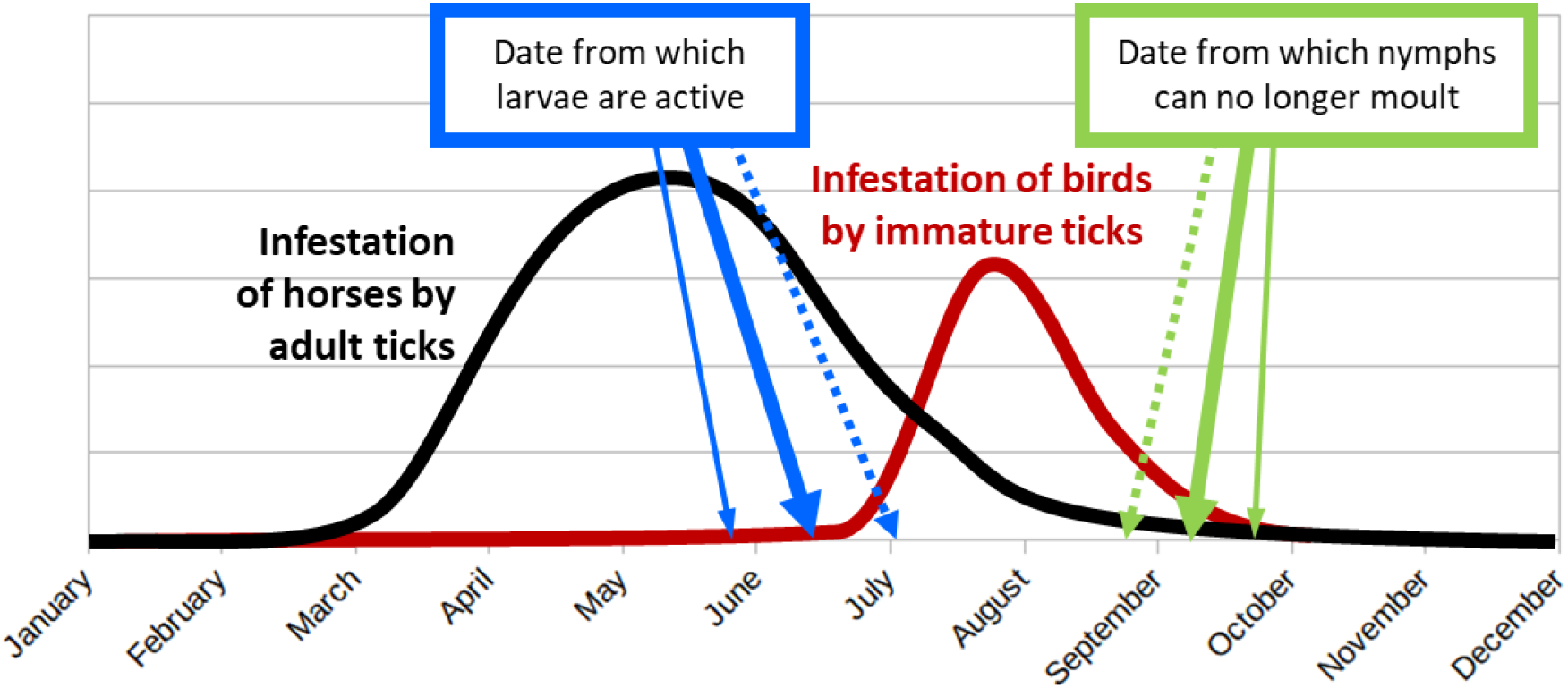
Stylized cycle of *Hyalomma marginatum* in the described study site of southern France. The thick plain arrows symbolize the present situation. The small plain arrows symbolize the potential situation with a warm spring and/or a mild autumn. The dotted arrows present the possible situation with a colder summer and/or an early autumn.

Tick infestation peaks in May. Afterwards, the population of available adult ticks in the environment gradually decreases as they find a host or die for different reasons, including exhaustion. Adult ticks can however resist this exhaustion for quite a long time if no host is available. The females, which engorge and drop off from March onwards, do not seem to lay eggs immediately, or the eggs do not hatch straight away. Larvae are active and can infest their hosts, in this case mainly ground-dwelling birds, from only the second half of June. Three weeks later, the first engorged nymphs are found in the environment. Quickly, in about 3-4 weeks, they moult into adults which remain inactive in the litter, undergoing a behavioural diapause. Immatures can engorge on birds until the end of September, but nymphs that detach too late are unable to moult before the onset of the cold season and most of them die during the winter. On the contrary, if engorged females detach late from their hosts, egg-laying may be delayed until the following spring.

The situation in southern France is therefore similar to that described in Turkey by Vatansever (Gray et al., 2009): “If temperatures are high enough to allow molting before the cold winters, unfed adults can survive the next active season. Field observations recorded the feeding of nymphs in late summer in Turkey, with the resulting unfed flat adults commonly overwintering in the first few centimetres below the soil surface.” Conversely, the phenology of *H. marginatum* may be slightly different in other regions of France where the tick is endemic. The species is now established in a vast area covering 300 km from south to north (42.50° N to 44.80° N) and 400 km from west to east (2.25° E to 6.90° E) (Bah et al., 2022). Therefore, the period of activity of ticks, both adults and larvae, could vary by a few weeks between these different zones, which should be further investigated.

Current climatic change could lead to warmer springs and milder autumns. In that case, larvae might become active earlier, while engorged nymphs might benefit from favourable moulting conditions for a longer period (small plain arrows of Figure 5). On the other hand, with a colder summer and/or an early autumn (dotted arrows in Figure 5), the larvae will be active too late and/or the nymphs will not be able to moult before the onset of the cold season. This is probably the situation in French regions where the climate is not Mediterranean and the tick is not established. This was also very likely the situation that prevailed in southern France until recently. At the end of the twentieth century, Morel (2003) pointed out that the French coast was the coolest part of the Mediterranean region which could explain why *H. marginatum* was not yet established in this area. Current global warming has probably corrected this “anomaly”. It is to be feared that, in the coming decades, the climate of other French regions will become favourable to this tick which will be able to establish in these new areas due to the movement of its hosts.

## Conclusion

In this study site of southern France, *H. marginatum* adults infested their host between March and August, with a peak in May. The larvae were active from mid- or end of June, being thus not yet available in spring to infest the migratory birds passing through the area. Engorged nymphs moulted in 3-4 weeks in summer but most died if the metamorphosis could not take place before the onset of cold conditions or gave adult ticks unable to extricate from the exuvia and deformed. Unfed adults that moulted in summer displayed a behavioural diapause until the following spring. Engorged females could survive the cold condition for up to 6 months before laying eggs. Predators were present in the environment but their impact was highly variable. To determine the exact kinetics of *H. marginatum* development (nymph moulting, oviposition and egg incubation) according to the temperature, experiments were carried out at the Baillarguet insectarium under controlled conditions. They are presented in other articles.

## Author contributions

AA identified the infested horse herd and participated in the horse survey between 2016 and 2018. FS organized and carried out the horse survey during the whole study period. MD, LV, and TP participated in the horse survey. GB planned the bird study and implemented it with FS. FS designed the experimental “cages” system and carried it out with MD. FS, LV and MD identified the collected ticks. CG carried out the molecular taxonomy analyses. FS analysed the data and wrote the first version of the manuscript, which was amended by MD. TP, LV, AA, CG and GB revised the manuscript. All authors approved the final version of the manuscript.

## Acknowledgements

The authors would like to thank the owner of the land where all the field studies were carried out. Several students occasionally helped to collect the ticks attached to the horses along the study years. Iyonna Zortman, a native English speaker, aided in correcting the language.

## Data and Supplementary Material availability

All data presented here and supplementary materials are available at the address https://doi.org/10.18167/DVN1/UKVEIU. The supplementary materials are:

S1. Meteorological data recorded by Météo-France at a site 13 km away from the study site.

S2. Infestation of the monitored horse herd by the ticks during the whole survey.

S3. Infestation of the examined birds by the ticks during the whole survey.

S4. Pictures of predated ticks.

S5. Pictures of normal and abnormal moulted adults.

The data available in the dataverse are:

D1. Data_Hyalomma_2016_2022.tab. Number of DNT for the whole study period. The data were used to plot Figure 1.

D2. Data_nymphs_cages_2019_2020.csv. Description of the nymphs and unfed adults recovered in the cages after release of engorged nymphs. The data were used to draw Figure 3.

D3. Data_females_cages_2019_2020.tab. Description of the ticks recovered in the cages after release of engorged females. The data were used to draw Figure 4.

## Footnotes

1 https://doi.org/10.18167/infrastructure/00001

2 Description of the weather station: https://www.meteociel.fr/obs/clim/meta/fiche_34274001.pdf. The monthly data can be downloaded from this website: https://www.meteociel.fr/climatologie/obs_villes.php?code2=34274001

## References

Apanaskevich DA, Horak I (2008) The genus Hyalomma Koch, 1844: V. re-evaluation of the taxonomic rank of taxa comprising the H. (Euhyalomma) marginatum koch complex of species (Acari: Ixodidae) with redescription of all parasitic stages and notes on biology. International Journal of Acarology, 34, 13–42. 10.1080/01647950808683704

Bacak E, Ozsemir AC, Akyildiz G, Gungor U, Bente D, Keles AG, Beskardes V, Kar S (2023) Bidirectional tick transport by migratory birds of the African-Western Palearctic flyway over Turkish Thrace: observation of the current situation and future projection. Parasitology Research, 123, 37. 10.1007/s00436-023-08069-x

Bah MT, Grosbois V, Stachurski F, Muñoz F, Duhayon M, Rakotoarivony I, Appelgren A, Calloix C, Noguera L, Mouillaud T, Andary C, Lancelot R, Huber K, Garros C, Leblond A, Vial L (2022) The Crimean-Congo haemorrhagic fever tick vector Hyalomma marginatum in the south of France: Modelling its distribution and determination of factors influencing its establishment in a newly invaded area. Transboundary and Emerging Diseases, n/a, 1–15. 10.1111/tbed.14578

Bakheit MA, Latif AA, Vatansever Z, U. S, Ahmed J (2012) The huge risks due to Hyalomma ticks. In: Arthropods as Vectors of Emerging Diseases (ed. Mehlhorn H), pp. 167–194. Springer, Berlin, Heidelberg (Germany). 10.1007/978-3-642-28842-5_8

Barré N, Mauléon H, Garris GI, Kermarrec A (1991) Predators of the tick Amblyomma variegatum (Acari: Ixodidae) in Guadeloupe, French West Indies. Experimental & Applied Acarology, 12, 163–170. 10.1007/BF01193464

Belozerov VN (1982) Diapause and biological rhythms in ticks. In: Physiology of ticks eds Obenchain FD and Galun R) (Vol. I), pp. 469–499. Pergamon Press, Oxford

Bernard C, Joly Kukla C, Rakotoarivony I, Duhayon M, Stachurski F, Huber K, Giupponi C, Zortman I, Holzmuller P, Pollet T, Jeanneau M, Mercey A, Vachiery N, Lefrançois T, Garros C, Michaud V, Comtet L, Despois L, Pourquier P, Picard C, Journeaux A, Thomas D, Godard S, Moissonnier E, Mely S, Sega M, Pannetier D, Baize S, Vial L (2024) Detection of Crimean–Congo haemorrhagic fever virus in Hyalomma marginatum ticks, southern France, May 2022 and April 2023. Eurosurveillance, 29, 2400023. 10.2807/1560-7917.ES.2024.29.6.2400023

Burns EC, Melancon DG (1977) Effect of imported fire ant (Hymenoptera: Formicidae) invasion on Lone star tick (Acarina: Ixodidae) populations. Journal of Medical Entomology, 14, 247–249. 10.1093/jmedent/14.2.247

Butler JF, Camino ML, Perez TO (1979) Boophilus microplus and the fire ant Solenopsis geminata. Recent Advances in Acarology, 1, 469–472

Capek M, Literak I, Kocianova E, Sychra O, Najer T, Trnka A, Kverek P (2014) Ticks of the Hyalomma marginatum complex transported by migratory birds into Central Europe. Ticks and Tickborne Diseases, 5, 489–493. 10.1016/j.ttbdis.2014.03.002

Chitimia-Dobler L, Schaper S, Rieß R, Bitterwolf K, Frangoulidis D, Bestehorn M, Springer A, Oehme R, Drehmann M, Lindau A, Mackenstedt U, Strube C, Dobler G (2019) Imported Hyalomma ticks in Germany in 2018. Parasites & Vectors, 12, 134. 10.1186/s13071-019-3380-4

De Liberato C, Frontoso R, Magliano A, Montemaggiori A, Autorino GL, Sala M, Bosworth A, Scicluna MT (2018) Monitoring for the possible introduction of Crimean-Congo haemorrhagic fever virus in Italy based on tick sampling on migratory birds and serological survey of sheep flocks. Preventive Veterinary Medicine, 149, 47–52. 10.1016/j.prevetmed.2017.10.014

Estrada-Peña A, Martínez Avilés M, Muñoz Reoyo MJ (2011) A population model to describe the distribution and seasonal dynamics of the tick Hyalomma marginatum in the mediterranean basin. Transboundary and Emerging Diseases, 58, 213–223. 10.1111/j.1865-1682.2010.01198.x

Gargili A, Thangamani S, Bente D (2013) Influence of laboratory animal hosts on the life cycle of Hyalomma marginatum and implications for an in vivo transmission model for Crimean-Congo hemorrhagic fever virus. Frontiers in cellular and infection microbiology, 3, 10. 10.3389/fcimb.2013.00039

Giupponi C, Jourdan-Pineau H, Bernard C, Blanda V, Bourquia M, Bru D, Cabezón O, Carrera-Faja L, Espunyes J, Gottlieb Y, Joly-Kukla C, Malandrin L, Mechouk N, Mihalca AD, Pollet T, Saengram P, Torina A, Valcárcel F, Vatansever Z, Vial L, Zahri A, Verheyden H, Huber K (2025) Tracking invasion events: phylogeography of Hyalomma marginatum in the Mediterranean basin with a focus on Southern France. Parasites & Vectors, 18, 407. 10.21203/rs.3.rs-5952236/v1

Grandi G, Chitimia-Dobler L, Choklikitumnuey P, Strube C, Springer A, Albihn A, Jaenson TGT, Omazic A (2020) First records of adult Hyalomma marginatum and H. rufipes ticks (Acari: Ixodidae) in Sweden. Ticks and Tick-borne Diseases, 11, 101403. 10.1016/j.ttbdis.2020.101403

Gray JS, Dautel H, Estrada-Peña A, Kahl O, Lindgren E (2009) Effects of Climate Change on Ticks and Tick-Borne Diseases in Europe. Interdisciplinary Perspectives on Infectious Diseases, 2009, 12 pages. doi:10.1155/2009/593232

Grech-Angelini S, Stachurski F, Lancelot R, Boissier J, Allienne J-F, Marco S, Maestrini O, Uilenberg G (2016) Ticks (Acari: Ixodidae) infesting cattle and some other domestic and wild hosts on the French Mediterranean island of Corsica. Parasites & Vectors, 9, 582. 10.1186/s13071-016-1876-8

Hoogstraal H, Kaiser MN, Traylor MA, Guindy E, Gaber S (1963) Ticks (Ixodidae) on birds migrating from Europe and Asia to Africa 1959-61. Bulletin of the World Health Organization, 28, 235–262

Hosseini-Chegeni A, Asadi M, Tavakoli M (2019) The record of Alveonasus canestrinii and Hyalomma marginatum (Acari:Ixodoidea) parasitizing partridge, Alectoris chukar (Aves: Phasianidae) in Iran. International Journal of Acarology, 45, 512–515. 10.1080/01647954.2019.1694981

Jaenson TGT, Tälleklint L, Lundqvist L, Olsen B, Chirico J, Mejlon H (1994) Geographical Distribution, Host Associations, and Vector Roles of Ticks (Acari: Ixodidae, Argasidae) in Sweden. Journal of Medical Entomology, 31, 240–256. 10.1093/jmedent/31.2.240

Jameson LJ, Morgan PJ, Medlock JM, Watola G, Vaux AGC (2012) Importation of Hyalomma marginatum, vector of Crimean-Congo haemorrhagic fever virus, into the United Kingdom by migratory birds. Ticks and Tick-borne Diseases, 3, 95–99. 10.1016/j.ttbdis.2011.12.002

Joly-Kukla C, Stachurski F, Duhayon M, Galon C, Moutailler S, Pollet T (2024) Temporal dynamics of the Hyalomma marginatum-borne pathogens in southern France. Current Research in Parasitology & Vector-Borne Diseases, 6, 100213. 10.1016/j.crpvbd.2024.100213

Lindeborg M, Barboutis C, Ehrenborg C, Fransson T, Jaenson TGT, Lindgren P-E, Lundkvist Å, Nyström F, Salaneck E, Waldenström J, Olsen B (2012) Migratory Birds, Ticks, and Crimean-Congo Hemorrhagic Fever Virus. Emerging Infectious Disease journal, 18, 2095. 10.3201/eid1812.120718

Morel P-C (1959) Les Hyalomma (Acariens, Ixodidae) de France. Annales de Parasitologie Humaine et Comparée, 34, 552–555

Morel P-C. (2003). Les tiques d’Afrique et du bassin méditerranéen. Distribution, Biologie, Ecologie, Rôle pathogène. In Manuscrit d’un livre jamais imprimé du fait du décès de l’auteur, publié sous forme de CD-Rom

Mwangi EN, Newson RM, Kaaya GP (1991) Predation of free-living engorged female Rhipicephalus appendiculatus. Experimental & Applied Acarology, 12, 153–162. 10.1007/bf01193463

Norval RAI, Rechav Y (1979) An assembly pheromone and its perception in the tick Amblyomma variegatum (Acarina : Ixodidae). Journal of Medical Entomology, 16, 507–511. 10.1093/jmedent/16.6.507

Nowak-Chmura M (2014) A biological/medical review of alien tick species (Acari: Ixodida) accidentally transferred to Poland. Annals of Parasitology, 60, 49–59

Palomar AM, Portillo A, Santibáñez P, Mazuelas D, Arizaga J, Crespo A, Gutiérrez Ó, Cuadrado JF, Oteo JA (2013) Crimean-Congo hemorrhagic fever virus in ticks from migratory birds, Morocco. Emerging Infectious Diseases, 19, 260–263. 10.3201/eid1902.121193

Pascucci I, Di Domenico M, Capobianco Dondona G, Di Gennaro A, Polci A, Capobianco Dondona A, Mancuso E, Cammà C, Savini G, Cecere JG, Spina F, Monaco F (2019) Assessing the role of migratory birds in the introduction of ticks and tick-borne pathogens from African countries: An Italian experience. Ticks and Tick-borne Diseases, 10, 101272. 10.1016/j.ttbdis.2019.101272

Pérez-Eid C (2007) Les Tiques. Identification, biologie, importance médicale et vétérinaire.

Monographies de microbiologie Lavoisier, Cachan, France Petney TN, Horak IG, Rechav Y (1987) The ecology of the african vectors of heartwater, with particular reference to Amblyomma hebraeum and A.variegatum. Onderstepoort Journal of veterinary Research, 54, 381–395

Portillo A, Palomar AM, Santibáñez P, Oteo JA (2021) Epidemiological Aspects of Crimean-Congo Hemorrhagic Fever in Western Europe: What about the Future? Microorganisms, 9. 10.3390/microorganisms9030649

Rahajarison P, Arimanana AH, Raliniaina M, Stachurski F (2014) Survival and moulting of Amblyomma variegatum nymphs under cold conditions of the Malagasy highlands. Infection, Genetics and Evolution, 28, 666–675. 10.1016/j.meegid.2014.06.022

Rechav Y, Parolis H, Whitehead GB, Knight MM (1977) Evidence for an Assembly Pheromone(S) Produced by Males of the Bont Tick, Amblyomma Hebraeum (Acarina: Ixodidae). Journal of Medical Entomology, 14, 71–78. 10.1093/jmedent/14.1.71

Samish M, Alekseev E (2001) Arthropods as predators of ticks. Journal of Medical Entomology, 38, 1–11. 10.1603/0022-2585-38.1.1

Santos-Silva MM, Vatansever Z (2017) Hyalomma marginatum Koch, 1844. In: Ticks of Europe and North Africa: A Guide to Species Identification eds Estrada-Peña A, Mihalca AD, and Petney TN), pp. 349–354. Springer International Publishing, Cham. 10.1007/978-3-319-63760-0_66

Sormunen JJ, Klemola T, Vesterinen EJ (2021) Ticks (Acari: Ixodidae) parasitizing migrating and local breeding birds in Finland. Experimental & Applied Acarology. 10.1007/s10493-021-00679-3

Stachurski F, Zoungrana S, Konkobo M (2010) Moulting and survival of Amblyomma variegatum (Acari: Ixodidae) nymphs in quasi-natural conditions in Burkina Faso; tick predators as an important limiting factor. Experimental & Applied Acarology, 52, 363–376. 10.1007/s10493-010-9370-z

Uiterwijk M, Ibáñez-Justicia A, van de Vossenberg B, Jacobs F, Overgaauw P, Nijsse R, Dabekaussen C, Stroo A, Sprong H (2021) Imported Hyalomma ticks in the Netherlands 2018–2020. Parasites & Vectors, 14, 244. 10.1186/s13071-021-04738-x

Vial L, Stachurski F, Leblond A, Huber K, Vourc’h G, René-Martellet M, Desjardins I, Balança G, Grosbois V, Pradier S, Gély M, Appelgren A, Estrada-Peña A (2016) Strong evidence for the presence of the tick Hyalomma marginatum Koch, 1844 in southern continental France. Ticks and Tick-borne Diseases, 7, 1162–1167. 10.1016/j.ttbdis.2016.08.002

Vinogradov DD, Smagin AS, Belova OA, Tsurikov SM, Karganova GG, Tiunov AV (2023) Identification of soil-dwelling predators of tick nymphs (Acari: Ixodidae) by stable isotope labeling. Journal of Medical Entomology. 10.1093/jme/tjad160

Walker AR, Bouattour A, Camicas J-L, Estrada-Peña A, Horak I, Latif AA, Pegram RG, Preston PM (2003) Ticks of domestic animals in Africa: a guide to identification of species. Bioscience Reports, Edinburgh

Weigand A, Teixeira J, Christian S (2020) First record of Hyalomma marginatum sensu stricto C.L. Koch, 1844 and distribution of Dermacentor reticulatus (Fabricius, 1794) (Acari, Ixodidae) in Luxembourg. Bulletin de la Société des naturalistes luxembourgeois, 122, 253–263

Wilkinson PR (1970) Factors affecting the distribution and abundance of the cattle tick in Australia: observations and hypotheses. Acarologia, XII, 492-508

Zingg S, Dolle P, Voordouw MJ, Kern M (2018) The negative effect of wood ant presence on tick abundance. Parasites & Vectors, 11, 164. 10.1186/s13071-018-2712-0

